# Cellular taxonomy of the preleukemic bone marrow niche of acute myeloid leukemia

**DOI:** 10.1101/2023.09.01.553259

**Authors:** Chinmayee Goda, Rohan Kulkarni, Alexander Rudich, Malith Karunasiri, Ozlen Balcioglu, Yzen Al-Marrawi, Erin Korn, Sanjay Kanna, Elizabeth A.R. Garfinkle, Bethany Mundy-Bosse, Bin Zhang, Guido Marcucci, Elaine R Mardis, Ramiro Garzon, Katherine E Miller, Adrienne M Dorrance

## Abstract

Mutations in hematopoietic stem/progenitor cells (HSPCs) can remain dormant within the bone marrow (BM) for decades before leukemia onset. Understanding the mechanisms by which these mutant clones eventually slead to full blown leukemia is of critical importance to develop strategies to eliminate these clones before they achieve their full leukemogenic potential. Recent data suggest that leukemic stem cells (LSCs) induce alterations within BM microenvironment (BMM) favoring LSC growth over normal HSCs. However, the cross talk between preleukemic stem cells (pLSC) and BMM is not completely understood. We hypothesize that pLSC induces critical changes within the BMM that are critical for leukemogenesis. To address this question, we are using our previously developed murine model of AML that highly recapitulates the human disease, develops AML sporadically with a preleukemic phase in which mice display normal white blood counts (WBCs) and absence of blasts in the BM. Thus, this is an excellent model to evaluate changes in the BMM that occurs during progression into AML. Using this model we performed single cell RNA-sequencing on cells from the BMM compared to wild-type (WT) controls. Overall, we defined the transcriptional profiles of pre-leukemic BMM cells and observed decreased percentages of normal BMM cells such as LepR+ mesenchymal stem cells (MSCs) and endothelial cells (ECs), known to regulate normal HSC function. Concomitantly, we found increases in CD55+ fibroblasts and NG2+ pericytes, that might play a more important role in regulation of pre-LSCs. Preleukemic CD55+ fibroblasts had a higher proliferation rate and showed significant down-regulation of several collagen genes known for regulating extra cellular matrix (ECM) including: *Col1a1, Col1a2, Col3a1, Col4a1*, and *Col6a1,* suggesting that ECM remodeling occurs in the early stages of leukemogenesis. Importantly, co-culture assays found that pre-leukemic CD55+ BM fibroblasts expanded pre-LSCs significantly over normal HSCs. In conclusion, we have identified distinct changes in the preleukemic BMM and identified a novel CD55+ fibroblast population that is expanded in preleukemic BMM that promote the fitness of pre-LSCs over normal HSCs.

**STATEMENT OF SIGNIFICANCE:** We have identified changes in the BMM landscape that define a preleukemic BM niche which includes the expansion of a novel CD55+ fibroblast population. These data suggest that a distinct preleukemic BM niche exists and preferentially supports LSC survival and expansion over normal HSCs to promote leukemogenesis.

## INTRODUCTION

Acute myeloid leukemia (AML) is a clonal, malignant disease of the blood and bone marrow (BM) that is heterogeneous at the molecular, cytogenetic, and clinical levels^1^. Non-random chromosomal abnormalities (e.g., deletions, translocations) are identified in 50-55% of all AML patients. In contrast, about 45-50% of all AML cases are cytogenetically normal (CN-AML)^2–5^. While advances have been made towards understanding the biology of AML, the prognosis is still very poor and there is an unmet need to develop novel and effective therapies^4,5^. Over the years it is becoming evident that along with genetic mutations within the hematopoietic stem and progenitor cell (HSPC) compartment, alterations in the BM microenvironment (BMM) also play an important role in transformation, therapy resistance and relapse^6^. Thus, to develop effective AML therapies, it is critical to understand how the intrinsic genetic alterations in HSPCs cooperate with the BMM to facilitate AML initiation, persistence, and therapy resistance. While alterations in BMM have been described in full blown leukemia, the BMM remodeling that happens during the preleukemic stage by HSCs with predisposing AML genetic mutations, have not been fully examined. Most studies on BM niche have utilized either transgenic or viral transduction murine models that induce non-physiologic expression of strong oncogenes that can cause aggressive disease by itself such as the viral transduction of HSPCs with *MLL(KMT2A)-*translocations, or PDX models that lack a preleukemic phase. Thus, investigating the changes that occur in the BMM before overt disease is present may help us to identify potential biomarkers or novel factors that contribute to preleukemic clonal expansion and ultimately to leukemia progression. These mechanisms have been difficult to identify previously since full blown disease results in massive blast infiltration of the BM, or they may no longer be required after the preleukemic stage.

Our lab has previously developed a murine model of CN-AML (*Mll*^PTD/WT^; *Flt3*^ITD/WT^, hereafter referred to as PTD;ITD) that recapitulates the human disease^7^. This model develops a sporadic leukemia with ∼100% penetrance (latency 4 to >18 months) with a preleukemic phase consisting of normal white blood cell counts (WBC), decreases in red blood cells (RBCs) and absence of AML blasts, thus allowing investigation of changes that occur before full-blown leukemic transformation. Here, we use this model to perform single cell RNA-sequencing and, we found decreases in LepR+ mesenchymal stromal cells (MSCs) and VE-cadherin+ (CD144+) endothelial cells (ECs) -specifically in arteriolar and arteriole ECs with no differences in sinusoidal ECs, along with increases in CD55+ fibroblasts. *In vitro* assays and microscopic analysis revealed increases in the proliferation of preleukemic CD55+ fibroblasts, suggesting active expansion and remodeling of the BMM by the preleukemic HSCs. Additionally, we also report altered mRNA and protein expression of collagen genes by the preleukemic CD55+ fibroblasts compared to WT. Finally, we observe that CD55+ fibroblasts can support preleukemic HSPC growth *in vitro.* Together, these data present the first analysis of the preleukemic BMM that differs from previously reported fully transformed AML BM niche^8^. This preleukemic BM niche demonstrated decreases in MSCs and ECs with an increase in an activated CD55+ fibroblast population that preferentially expands in preleukemic HSCs.

## RESULTS

### Global alterations in pre-leukemic bone marrow niche populations

To determine a relevant time-point to examine changes in PTD; ITD mice that would indicate BM dysfunction, WBC and RBC counts in peripheral blood were monitored every 2 weeks starting from 8 weeks after birth. Preleukemic stage was determined when we observed first consistent changes in peripheral blood (PB) cells without observing blasts in PB and bone marrow. For this murine model, the first hematology alteration we noticed was a decrease in RBC counts. Thus, preleukemic stage was defined when RBC counts were consistently low (RBC counts of less than 6 million/uL in 3 consecutive bleeds), without an elevation in WBC counts and presence of blasts in blood and BM (Supplemental Figure S1A). To profile the bone marrow niche in preleukemic mice, non-hematopoietic bone marrow stromal cells from WT and preleukemic PTD; ITD mice (n=3) were sorted by flow cytometry using the markers CD45-Gr1-CD11b-B220-CD3-CD19-Ter119-CD71- (Figure 1A, Supplemental Figure S1B) as previously described and analyzed by droplet-based single cell RNA-sequencing^8^. Data was pre-filtered based on RNA counts and removal of hematopoietic cells, resulting in a pool of 12,444 cells (WT = 5,677 cells, PTD; ITD = 6,767 cells) used for further analysis. Median number of genes detected per cell were 1531 for WT, 1394 for PTD; ITD and median number of transcripts per cell were 3206 for WT and 2998 for PTD; ITD. WT and preleukemic PTD; ITD stromal cells were integrated, and unbiased clustering was performed resulting in 17 clusters (Figure 1B). Cell types were annotated based on expression of cell-specific genes (Figure 1C-D, Supplemental Table 1) as previously described^8^. We identified MSCs (clusters 0, 1, 2, 3, 8, expressing *Lepr*, and *Cxcl12*), endothelial cells (clusters 4, 14, 18, expressing *Cdh5*, and *Pecam1*), fibroblasts (clusters 5,6, 10, 12 expressing *S100a4*), pericytes (cluster 7 expressing *Atp1b2*), osteo-lineage cells (OLCs, clusters 9, 13, 16 expressing *Bglap*), and chondrocytes (cluster 11, expressing *Acan*). We then analyzed the percentages of cells across clusters and between the samples.

**Figure 1.**
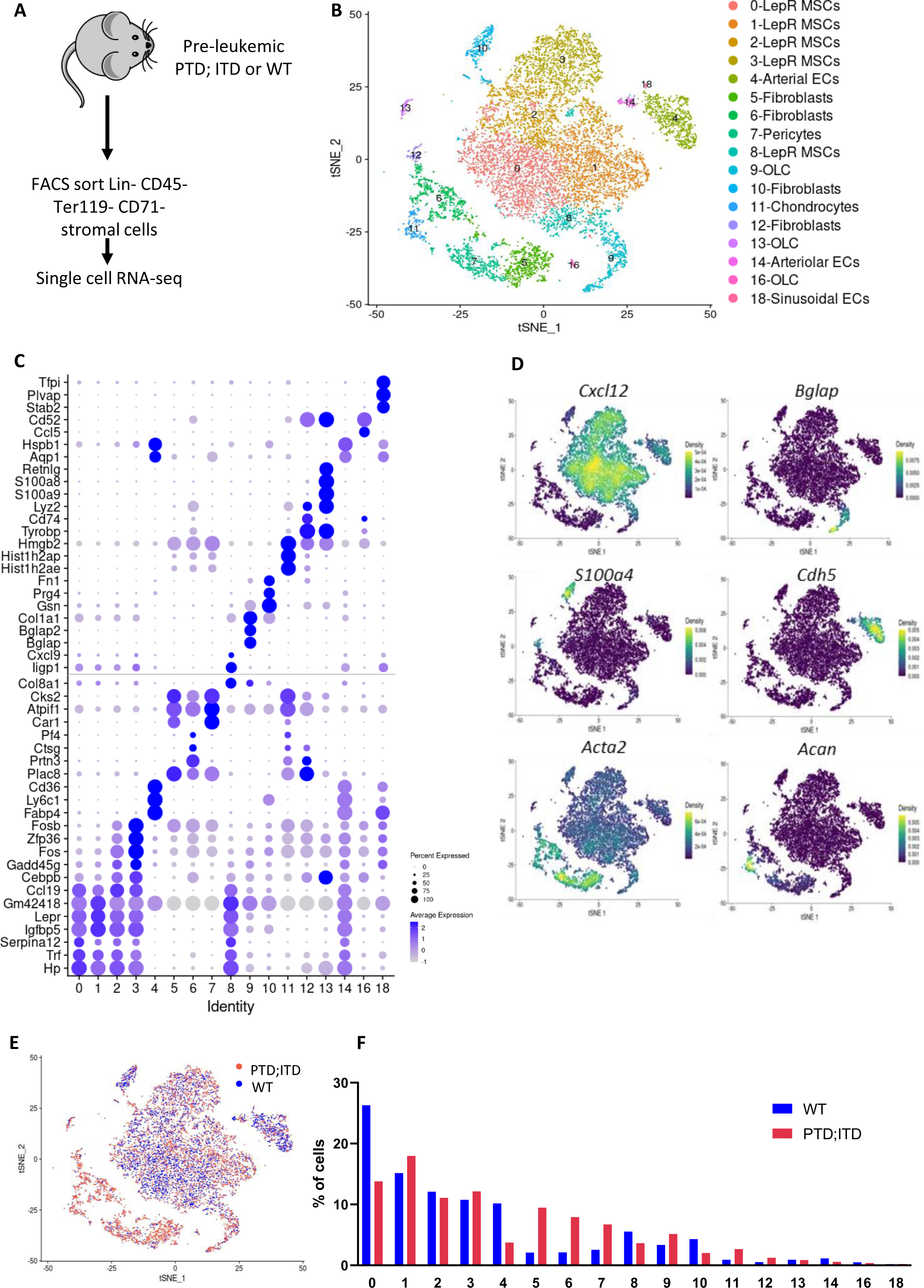
An atlas of preleukemic bone marrow stroma. **(A)** Schematic representation of experiment. Preleukemic PTD; ITD or age- and sex-matched WT mice were sacrificed. Femurs and tibias were collagenase digested and Lin-CD45-Ter119-CD71-stromal cells were sorted by flow cytometry. Sorted stromal cells were analyzed by 10x scRNA-sequencing. **(B)** t-Distributed stochastic neighbor embedding (t-SNE) of stromal cells, colored by clustering and annotated based on gene signature. (C) Dot plot of top expressed genes in stromal cells by clusters. **(D)** t-SNE of stromal cells by expression of key cell-type marker genes **(E)** t-SNE of stromal cells colored by condition (WT, blue; preleukemic PTD; ITD, red). **(F)** Percentage of stromal cells across each cluster in WT and preleukemic PTD; ITD cells.

We found decreases in percentages of LepR MSCs (cluster 0) and arterials ECs (cluster 4) while an increase in percentages of fibroblasts (clusters 5, 6, 12) and pericytes (cluster 7) in preleukemic BMM compared with their controls (Figures 1E-F). Furthermore, we analyzed the proliferation status of stromal cell subpopulations using the cell cycle scoring function included in Seurat package. Overall, we show decreases in proliferation of stromal cells (Supplemental Figure S2A) with decreases in the percentage of cycling cells (cells in S and G2M phase) in preleukemic LepR+ MSC and EC clusters with an increase in preleukemic fibroblasts compared to WT (Supplemental Figure S2B). These data demonstrate substantial changes in the preleukemic BMM that are distinct from WT.

### LepR MSC clusters 0 and 8 are lost in the BMM of pre-leukemic mice

LepR MSCs are multipotent stromal cells that have the ability to differentiate into osteo-lineage cells, adipocytes, and chondrocytes^12^. They also play a crucial role in regulating HSCs by secreting critical niche factors including Cxcl12, and Kit ligand^13^. Several studies have shown diverse roles of LepR MSCs in AML, either promoting or inhibiting AML cell proliferation through induction of proliferation or apoptosis respectively^14^. Since they play an important role in hematopoiesis and AML, we analyzed their presence within the pre-leukemic BM niche.

We identified 5 clusters of LepR MSCs - clusters 0, 1, 2, 3, and 8, expressing canonical LepR MSC markers^8^: *Lepr*, *Cxcl12*, *Kitl*, (Figure 2A-C). These cells comprised the majority of the stromal cells in normal as well as pre-leukemic BM. Consistent with their role in HSC regulation, all LepR MSC clusters expressed HSC niche factors *Cxcl12*, *Ccl2*, and *Cxcl14*, *Ccl19* (Figure 2D). First, we determined whether there were any differences in the percentages of LepR+ MSC clusters in preleukemic PTD; ITD bone marrow compared with its control. We found significant decreases in the cell percentages only in subclusters 0 (∼50%) and 8 (∼40%) LepR+ MSCs in preleukemic PTD; ITD BM compared to control (Figure 1F). Since we identified a decrease in LepR MSC clusters 0, and 8 in pre-leukemic BM, we further defined these LepR MSC subpopulations by analyzing the markers expressed by all cells in these clusters. Upon analysis of top markers expressed in LepR MSC cluster 0 (Supplemental Figure S3A), we identified these cells express *Nme2, Lgals1*, which have been previously shown to be important for anti-inflammatory response^15^, and *S100a6,* which is required for HSCs self-renewal and regeneration^16^. Similarly, we defined subcluster 8 and found that all cells in this cluster express osteo-lineage progenitor cell markers *Sp7*, and *Alpl*, which ultimately give rise to osteo-lineage cells, that play a crucial role in HSC maintenance^8^. Since LepR MSC clusters 0 and 8 are lost in pre-leukemic PTD;ITD mice, these data suggests that the subset of LepR MSCs required for maintenance and self-renewal of normal HSCs is lost within the pre-leukemic BM niche.

**Figure 2.**
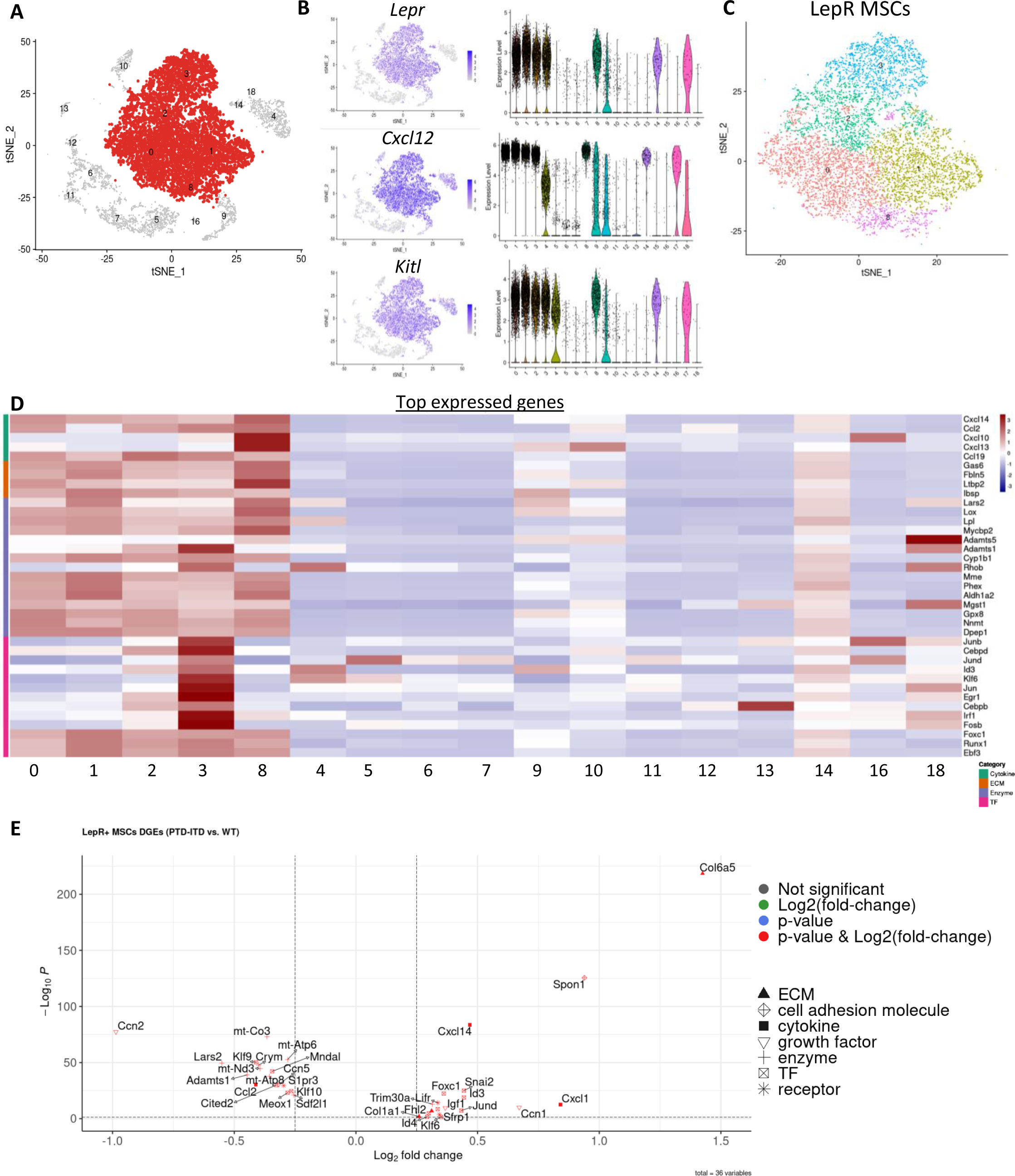
LepR+ MSCs in preleukemic bone marrow. **(A)** t-SNE of stromal cells, highlighting LepR MSC clusters. **(B)** t-SNE of stromal cells by expression of key MSC marker genes, along with the corresponding violin plot distributions of expression levels across clusters. **(C)** t-SNE of LepR MSC clusters. **(D)** Heatmap of top expressed genes (rows) in LepR MSC clusters in the cells of each cluster (columns) ordered by gene categories. **(E)** Differentially expressed genes in preleukemic PTD; ITD LepR clusters compared to WT.

Finally, to understand transcriptomic changes within all LepR MSCs in pre-leukemic niche, we analyzed genes differentially expressed in all LepR MSC clusters in pre-leukemic BM compared to normal. Gene expression analysis revealed increases in the expression of genes important for cell adhesion-*Spon1* (∼2-fold increase, *P<0.01*), cytokines and growth factors-*Ccn1*, (∼1.5-fold increase, *P<0.01*) *Cxcl14*, (∼2-fold increase, *P<0.01*), and *Cxcl1* (∼1.2-fold increase, *P<0.01*), and transcription factors-*Foxc1* (∼2.8-fold increase, *P<0.01*), *Snai2* (∼2.3-fold increase, *P<0.01*), *Id3* (∼2.3-fold increase, *P<0.01*), *Klf6* (∼2.9-fold increase, *P<0.01*), and *Jund* (∼2.3-fold increase, *P<0.01*) (Figure 2E). These data together indicate substantial changes in the cluster cell percentages and gene expression in the pre-leukemic LepR MSCs compared to normal BMM.

### Gain in extra cellular matrix (ECM)-secreting osteo-lineage cells (OLCs) in the pre-leukemic niche

MSCs can differentiate into osteo-lineage cells that comprise of osteoprogenitors and mature osteoblasts. Osteoblasts play a critical role in HSC maintenance, and differentiation and have been implicated in many diseases such as multiple myeloma and AML^17,18^. Studies have reported that the leukemic BM niche is marked by deficiency in osteogenesis and loss of mineralized bone, that ultimately disrupts the normal endosteal physiology. Given the importance of osteoblasts in hematopoiesis and leukemogenesis, we investigated the osteo-lineage cells in pre-leukemic BM.

We identified subclusters 9, 13, and 16 as osteo-lineage cells (OLCs), that express the definitive osteo-lineage markers^8^: *Runx2*, *Bglap*, and *Sp7* (Supplemental Figure S4A-B), secreted cytokines (*Ccl4*, *Ccl5*, *Ccl6*, and *Cxcl10*) and growth factors (*Bmp3*) (Supplemental Figure S4C). We analyzed whether there was lineage skewing in OLC population percentages in preleukemic PTD; ITD bone marrow compared to WT and found only an increased percentage of cluster 9 in preleukemic PTD; ITD stroma (∼1.5-fold increase) (Figure 1F). To define each cluster and understand the importance of OLC cluster 9 within the pre-leukemic niche, we identified the top expressed genes in each subcluster of OLCs in WT and pre-leukemic PTD;ITD BM and categorized them into secreted factors, extracellular matrix (ECM) genes, enzymes, and transcription factors. We found that cluster 9 had increased expression of ECM collagen genes (*Col1a1*, *Col2a1*, *Col5a2*. *Col11a1*, *Col11a2*, and *Col12a1*) compared to other OLC clusters (Supplemental Figure S4C), indicating that there is a gain in ECM-secreting OLCs in the pre-leukemic niche. This suggests remodeled ECM within pre-leukemic niche might be important for leukemogenesis.

### Vascular remodeling of pre-leukemic niches is different than in full-blown leukemias

BM endothelial cells (ECs) make up the vasculature within the BM that forms an essential component of the BMM. While the sinusoidal vessels act as a site for differentiation of certain hematopoietic progenitors, and as a conduit for mature hematopoietic cells to peripheral circulation, the endosteal arteries and arterioles are important for maintaining HSC quiescence^19^. In the BM of AML patients, BM microvascular density increases, which ultimately promotes the proliferation and mobilization of AML blasts into circulation^20^. We wanted to analyze whether vascular remodeling occurred within the pre-leukemic BM and characterize these changes.

Consistent with previous studies, we identified 3 EC clusters: 4, 14, and 18, based on the expression of established EC marker genes: *Cdh5*, *Pecam1*, and *Emcn* (Figure 3A). To determine the subtypes of ECs, we analyzed expression of *Vwf, Ly6a*^high^ for arterial (cluster 4), *LepR* for arteriolar (cluster 14), and *Flt4* for sinusoidal (cluster 18) ECs as previously described (Figure 3B-C, Supplemental Figure S5)^8^. To determine changes in pre-leukemic BM vasculature, we first analyzed the percentage of cells in each EC cluster in preleukemic PTD; ITD and WT BM. Contrary to what is observed in the BM of full-blown AML, we found a ∼2.8 decrease in arterial ECs (cluster 4) and ∼2-fold decrease in arteriolar ECs (cluster 14), with no difference in the percentage of sinusoidal ECs (cluster 18) in preleukemic PTD; ITD bone marrow compared to WT (Figure 1F). Finally, we identified differentially expressed genes in preleukemic PTD; ITD EC clusters compared to WT. We identified chemokine *Cxcl9* was upregulated in PTD;ITD ECs compared to WT (∼1.4-fold increase, *P<0.01*), which has been shown to be induce chemotaxis, and disrupt endothelial barrier function previously^21^. *Ccn2* (∼ 2-fold decrease, *P<0.01*), known to regulate endothelial cell adhesion to pericytes^22^, and *Cacybp* (∼1.3-fold decrease, *P<0.01*), previously shown inhibit vascular remodeling^23^, were decreased in preleukemic PTD; ITD ECs compared to WT ECs (Figure 3D), confirming vascular remodeling of BM niche in pre-leukemic PTD;ITD BM.

**Figure 3.**
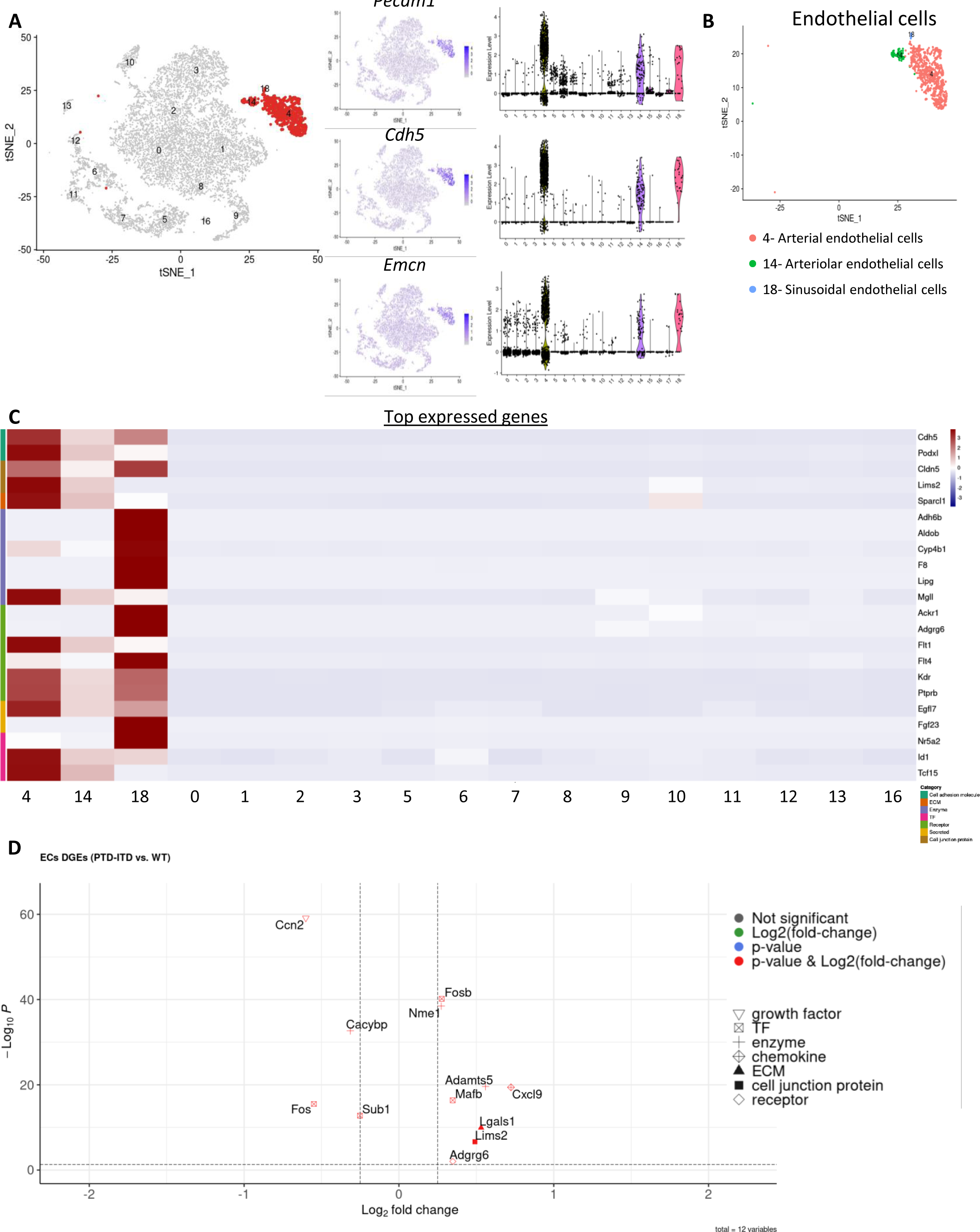
Endothelial cells in preleukemic bone marrow. **(A)** t-SNE of non-hematopoietic cells, and expression of key endothelial cell marker genes, along with the corresponding distributions of expression levels across clusters. **(B)** t-SNE of three subclusters in the endothelial cell cluster. **(C)** Expression of top expressed genes (rows) in the cells of each cluster (columns) ordered by gene categories. **(D)** Differentially expressed genes in preleukemic PTD; ITD EC clusters compared to WT.

### Pericytes are increased in pre-leukemic bone marrow niche

Pericytes are mural cells that are typically found along ECs, and in addition to providing structural support to vasculature also play an important role in regulating hematopoiesis ^24^. While pericytes have not been described in detail in AML, preliminary studies have shown a decrease in pericyte coverage of BM vasculature in bone marrow biopsies of AML patients^25^. To determine whether pericytes are altered in the pre-leukemic niche, we analyzed the pericytes population in our dataset.

We identified cluster 7 as pericytes based on *Atp1b2* expression (Figure 4A-B). Upon analysis of top markers expressed by all pericytes, we found cell adhesion markers *Ninj1*, enzymes *Cdk6*, *Khk*, and *Gclm* and transcription factors *Ikzf1*, *Gata1*, and *Hdgf* expressed in pericytes from WT and pre-leukemic BM (Figure 4C). To determine if the perivascular niche is changed in pre-leukemic PTD;ITD BM, we first analyzed the percentage of pericytes in pre-leukemic BM compared to WT. Contrary to loss of pericytes in AML BM niche^26^, we observed a ∼2.7-fold increase in percentage of pericytes in preleukemic PTD; ITD BM compared to WT BM (Figure 1F), suggesting a unique perivascular niche during leukemogenesis. Finally, to understand the transcriptomic changes in pre-leukemic pericytes, we identified the differentially expressed genes in pericytes in PTD; ITD bone marrow compared to WT, and categorized them into transcription factors, enzymes, and transmembrane signal receptors. Transcription factors *Fosb* (∼2.6-fold increase, *P<0.01*), *Klf2* (∼3.4-fold increase, *P<0.01*), and *Klf6* (∼2.2-fold increase, *P<0.01*), known to regulate pericyte recruitment^27^ were increased in preleukemic PTD; ITD bone marrow compared to WT (Figure 4D), confirming an altered perivascular niche in pre-leukemic BM.

**Figure 4.**
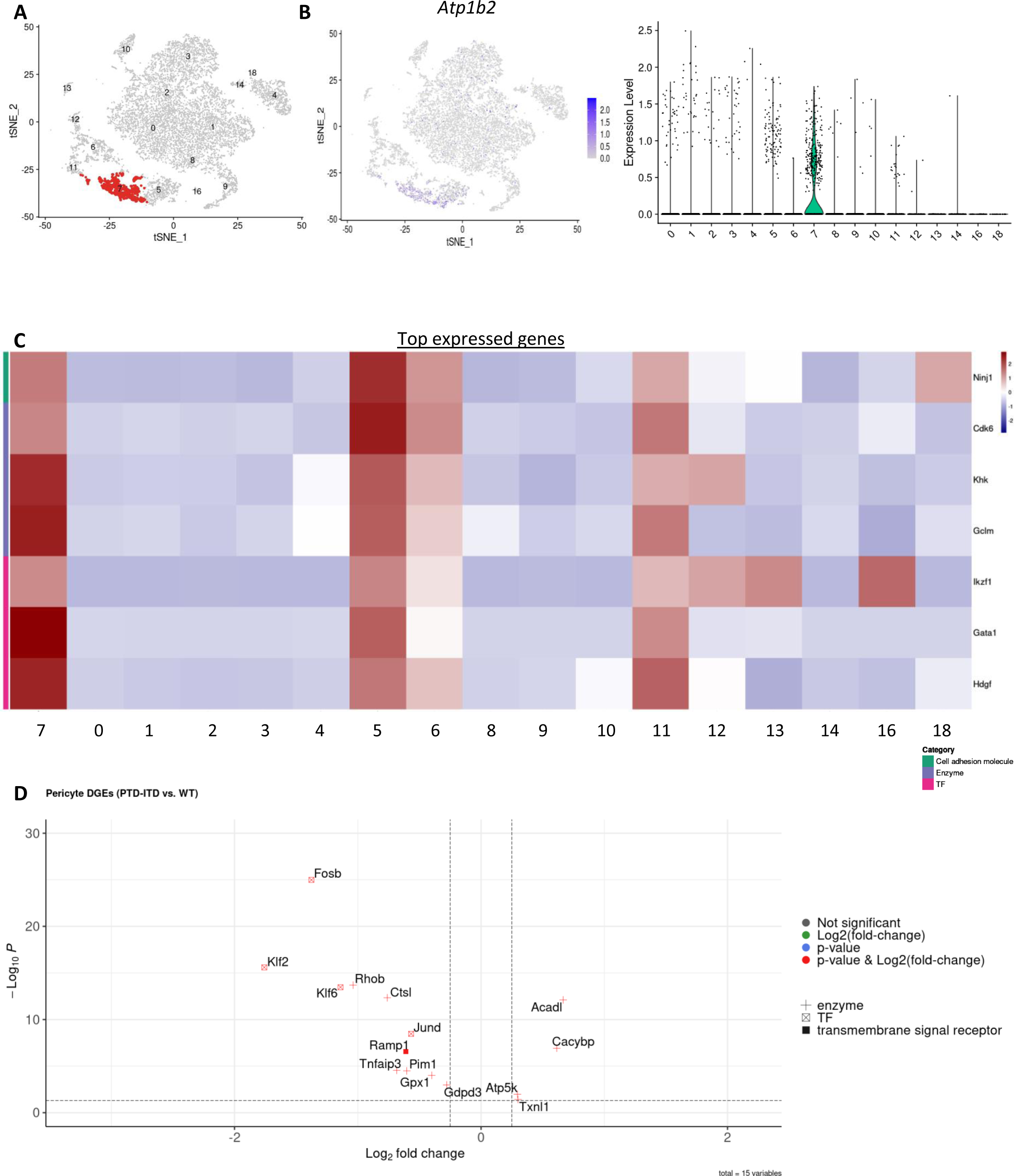
Pericytes in preleukemic bone marrow. **(A)** t-SNE of stromal cells, highlighting pericyte cluster. **(B)** t-SNE of stromal cells by expression of key pericyte marker gene, along with the corresponding distributions of expression level across clusters. **(C)** Heatmap of average expression of top expressed genes (rows) of pericyte cluster in the cells of each cluster (columns) ordered by gene categories. **(D)** Volcano plot of differentially expressed genes in preleukemic PTD; ITD pericyte cluster compared to WT.

### Bone marrow CD-55+ fibroblasts are increased in pre-leukemic niche

Bone marrow fibroblasts are stromal cells that provide mechanical support to HSPCs by secreting extracellular matrix and remodeling of extracellular matrix components into a highly structured network, thereby regulating hematopoiesis^28^. Several studies using human bone marrow fibroblasts have shown that they can protect AML blasts from undergoing apoptosis and cause resistance to therapy *in vitro*^29,30^. Given their importance in AML pathogenesis and response to therapy, we analyzed the BM fibroblasts in the pre-leukemic niche.

We identified 4 fibroblast subclusters: 5, 6, 10, and 12, based on the expression of the fibroblast marker genes: *S100a4*, *Sema3c*, and *Dcn* (Figure 5A-B). To determine the changes in fibroblast populations within the pre-leukemic BM, we first analyzed population differences compared to WT BM. Interestingly, we found an increase in fibroblast clusters 5 (∼4.8-fold increase), 6 (∼3.7-fold increase), and 12 (∼2.5-fold increase), while a decrease in fibroblast cluster 10 (∼2-fold decrease). To better understand the different fibroblast clusters, we decided to characterize them based on gene expression. All cells in fibroblast clusters 6, and 12 expresses *Spi1*, a gene previously shown to be an essential regulator of pro-fibrotic gene expression program in fibroblasts^31^ (Supplemental Figure S6A). Cluster 10, which is lost in the pre-leukemic BM, showed elevated expressions of ECM genes which includes: *Col3a1*, *Col6a3*, *Fbln2*, *Lgals1*, and *Tnxb* (Figure 5C). This suggests that ECM secreting fibroblasts modulate the pre-leukemic ECM. Additionally, upon analyzing the differentially expressed genes in preleukemic PTD; ITD fibroblast clusters, we found decreased expression of multiple collagen genes-*Col1a1* (∼4.8-fold decrease, *P<0.01*), *Col1a2* (∼3.6-fold decrease, *P<0.01*), *Col3a1* (∼4.3-fold decrease, *P<0.01*), *Col4a1* (∼1.8-fold decrease, *P<0.01*), *Col5a2* (∼2.5-fold decrease, *P<0.01*), in preleukemic PTD; ITD fibroblasts compared to WT (Figure 5D), confirming a loss of ECM in pre-leukemic niche.

**Figure 5.**
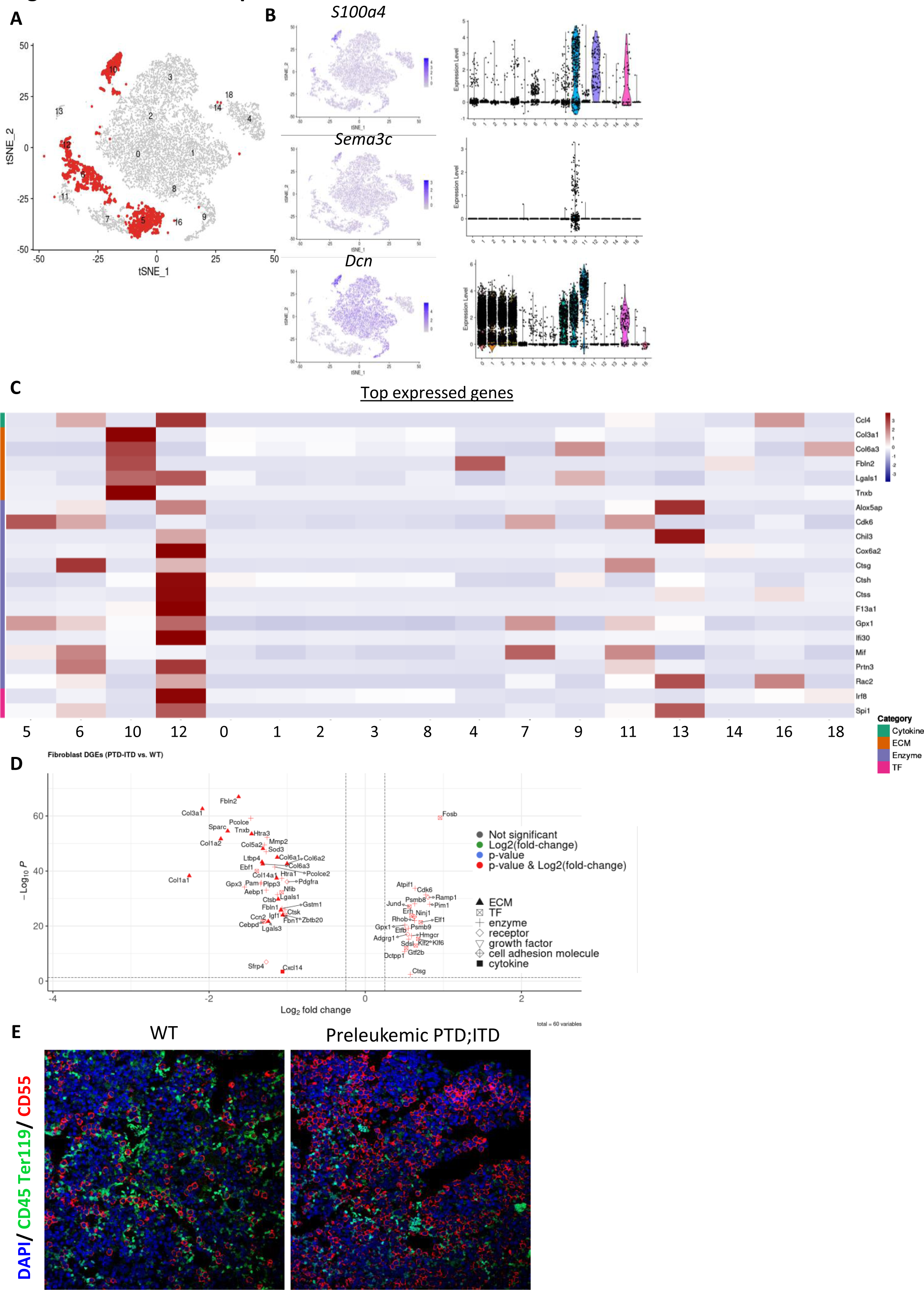
Fibroblasts in preleukemic bone marrow. **(A)** t-SNE of stromal cells, highlighting fibroblast clusters. **(B)** t-SNE of stromal cells by expression of key fibroblast marker genes, along with the corresponding violin plot distributions of expression levels across clusters. **(C)** Four subclusters in the fibroblasts cluster. **(D)** Heatmap of expression of top expressed genes (rows) of fibroblasts clusters in the cells of each cluster (columns) ordered by gene categories. **(E)** Volcano plot of differentially expressed genes in preleukemic PTD; ITD fibroblast clusters compared to WT. **(F)** Representative immunofluorescence staining for CD45 (green), Ter119 (green), and CD55 (red) in bone sections from WT and preleukemic mice. Nuclei are counterstained with DAPI. Images are representative of N = 3 per group.

Since we observed an overall increase in BM fibroblasts within the pre-leukemic niche, we analyzed expression of cell surface markers in order to investigate them further. We identified *Cd44*, and *Cd55* to be up regulated in preleukemic PTD; ITD (Supplemental Figure S6B). We validated this finding by performing immunofluorescence on femur sections from preleukemic PTD; ITD and WT mice. We observe an increase in CD45-Ter119-CD55+ fibroblasts in bone marrow of preleukemic PTD; ITD mice compared to WT (Figure 5E). Hence, we identified an expansion of CD55+ fibroblasts within the pre-leukemic BM.

To further characterize the CD55+ fibroblasts and understand their possible role in leukemogenesis, we first sorted CD55+ stromal cells from WT and pre-leukemic PTD;ITD mice and performed scRNA-seq. Upon clustering, we identified 6 sub-clusters within CD55+ fibroblasts (Supplemental Figure S7A). Upon analysis of distribution of cells across clusters, we found a loss of cluster 2 and increase in clusters 3, 4, and 5 in PTD;ITD CD55+ fibroblasts compared to WT fibroblasts (Supplemental Figures S7B-C). Further analysis identified that CD55+ fibroblast cluster 2 had decreased proliferation score compared to other clusters, suggesting that there is a loss of non-cycling CD55+ fibroblasts in pre-leukemic PTD;ITD BM (Supplemental Figure S7D). To validate the scRNA-seq findings, we isolated CD55+ fibroblasts from WT and preleukemic PTD;ITD BM and cultured them *in vitro*. Preleukemic CD55+ fibroblasts showed increased growth *in vitro* compared to WT CD55+ fibroblasts as analyzed by MTS assay (Figure 6A). Furthermore, upon analysis of proliferation marker Ki67 by immunostaining CD55+ fibroblasts in BM, we observed an increased percentage of Ki67+ CD55+ fibroblasts in preleukemic BM (Figure 6B) confirming increased proliferation of CD55+ fibroblasts in preleukemic niche. These results support the increased percentage of CD55+ fibroblasts identified in the scRNA-seq data. We next validated the changes in extracellular matrix (ECM) proteins in preleukemic BM by performing immunofluorescence on femur sections. Consistent with the scRNA-seq data, we observed a decrease in collagen I (Figure 6C) and collagen IV (Figure 6D) staining in preleukemic BM compared to WT BM, confirming the scRNA-seq data. These results suggest that the preleukemic BM undergoes ECM remodeling, which may support expansion of LSCs. To investigate the effect of CD55+ fibroblasts on LSCs, we performed co-culture assays followed by CFU-C. cKit cells from WT or preleukemic PTD;ITD mice were co-cultured with CD55+ fibroblasts from WT or preleukemic mice followed by CFU-C assay. We observed an increase in CFU-C colonies when cKit+ cells from PTD;ITD mice were co-cultured with CD55+ fibroblasts from PTD;ITD mice compared to CD55+ fibroblasts from WT mice (Figure 6E). These results suggest that CD55+ fibroblasts expand within the preleukemic BM and support expansion of LSCs, thereby enabling leukemic progression.

**Figure 6.**
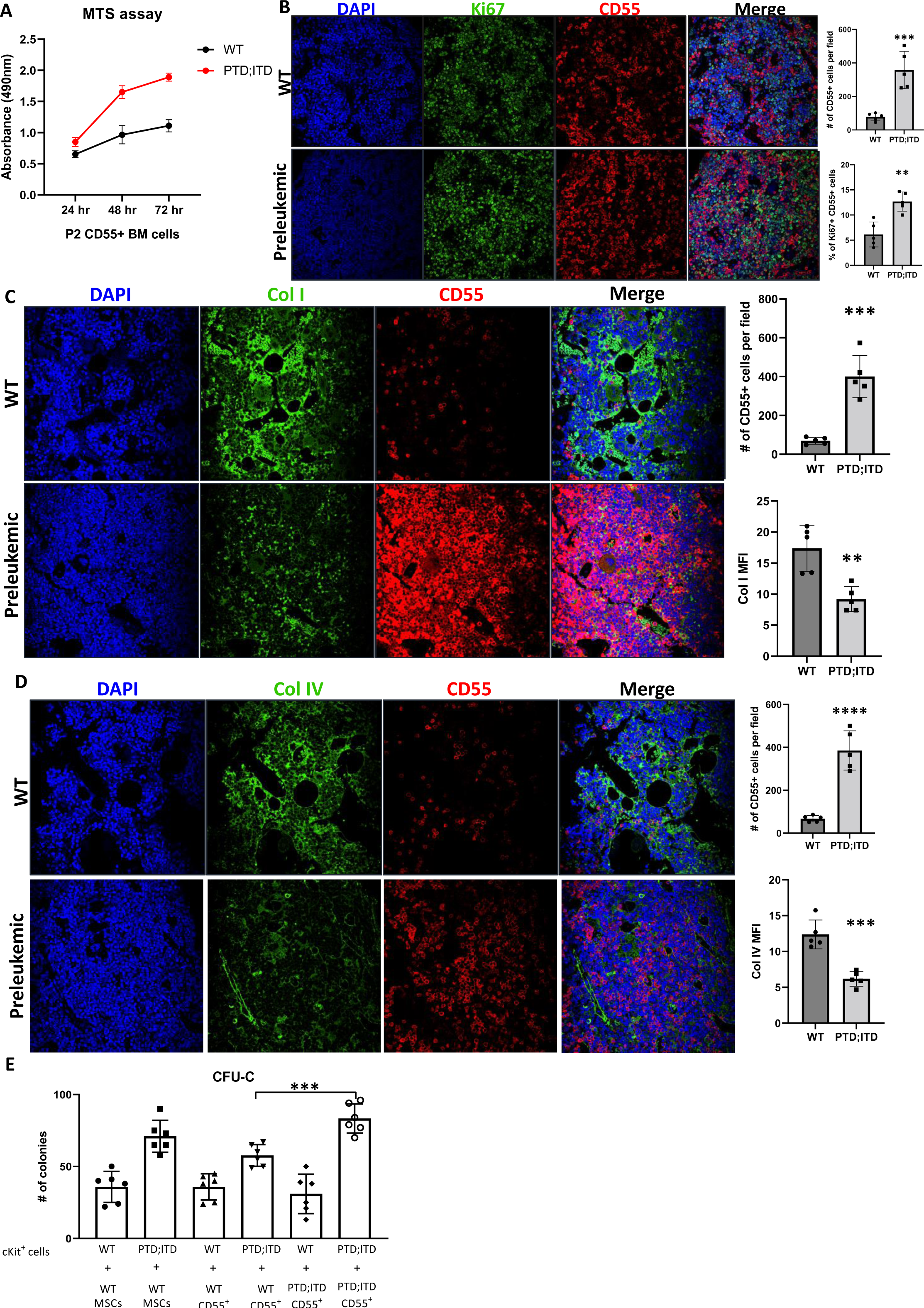
Expansion of CD55+ fibroblasts in preleukemic BM. **(A)** Growth analysis by MTS assay of P2 CD55+ fibroblasts from WT and preleukemic PTD;ITD mice at 24, 48, and 72 hours. N= 4 per group. **(B)** Representative immunofluorescence staining for Ki67 (green), and CD55 (red) in bone sections from WT and preleukemic PTD;ITD mice. Nuclei are counterstained with DAPI. Images are representative of N = 5 mice per group. (Right, top) Quantification of number of CD55+ cells per field in WT and preleukemic BM. (Right, bottom) Quantification of percentage of Ki67+ CD55+ cells per field in WT and preleukemic BM. **(C)** Representative immunofluorescence staining for Collagen I (green), and CD55 (red) in bone sections from WT and preleukemic PTD;ITD mice. Nuclei are counterstained with DAPI. Images are representative of N = 5 mice per group. (Right, top) Quantification of number of CD55+ cells per field in WT and preleukemic BM. (Right, bottom) Quantification of mean fluorescence intensity (MFI) of Collagen I staining per field in WT and preleukemic BM. **(D)** Representative immunofluorescence staining for Collagen IV (green), and CD55 (red) in bone sections from WT and preleukemic PTD;ITD mice. Nuclei are counterstained with DAPI. Images are representative of N = 5 mice per group. (Right, top) Quantification of number of CD55+ cells per field in WT and preleukemic BM. (right, bottom) Quantification of mean fluorescence intensity (MFI) of Collagen IV staining per field in WT and preleukemic BM. **(E)** Co-culture followed by CFU-C assay. Briefly, cKit+ cells from WT or pre-leukemic PTD;ITD mice were co-cultured with P2 WT MSCs or P2 CD55+ fibroblasts from WT or pre-leukemic PTD;ITD mice for 48 hours followed by CFU-C assay. Colonies were counted 14 days later. Quantification of number of colonies. N = 6 per group.

## DISCUSSION

The BM niche has been well characterized under homeostatic conditions and after the onset of leukemia^8^. However, the changes that take place in the BM niche prior to leukemic transformation has not been well studied. This is the first study to extensively characterize the BM niche cellular heterogeneity prior to leukemic transformation, thus allowing us to identify populations that might possibly be involved in promoting leukemic transformation. The PTD;ITD mouse model with a single copy of each mutant allele, has a prolonged latency period, which allows us to investigate changes that occur in the BM niche prior to leukemic transformation. Furthermore, this mouse model harbors both mutations under the control of their respective normal endogenous promoter thereby allowing physiological and temporal expression of mutant alleles. Using this model, we identified 17 stromal cell clusters with differential frequencies and gene expression patterns in the preleukemic BM niche compared to WT.

RBC counts and conditions of anemia have been frequently associated with leukemic transformation. Presence of moderate or severe anemia was identified as a predictor of leukemic transformation in patients with primary myelofibrosis^32^. Furthermore, previous study has shown that a decrease in erythroblasts and mature RBCs precedes leukemic transformation in a xenograft mouse model^33^. Hence to analyze the changes that take place in BM niche prior to leukemic transformation, we performed scRNA-sequencing on BM stromal cells at the time point when PTD; ITD mice had decreased peripheral blood RBC counts, without an increase in peripheral blood WBC counts.

The BM niche has been characterized in AML patients and in mouse models of AML after the establishment of the disease. Particularly, the anatomical and functional changes in leukemic BM vasculature have been previously characterized. Using PDX model of AML, a significant loss of ECs associated with sinusoids (CD31+Sca1low) as well as an increased number of ECs associated with arterioles (CD31+Sca1high) is observed in leukemic BM^26^. In our study, while we observe changes in gene expression patterns in EC clusters, we identify no observable differences in percentage of BM sinusoidal ECs. We see a decrease in the percentage of arterial and arteriolar ECs, subset of ECs known to secrete higher levels of niche factors supporting normal HSCs. Our data shows a loss of EC populations known to support normal HSCs in the pre-leukemic niche, suggesting an alteration of the BM vascular niche to support pre-leukemic HSPCs instead. Our results suggest that changes in BM EC numbers occur as a result of leukemic transformation and might not be required for leukemic transformation.

While frequencies of majority of BM stromal cell populations remained unchanged in our data, we observed a striking increase in pericyte and fibroblast populations in preleukemic BM compared to WT. BM fibroblasts have been characterized to some extent in patients with myelofibrosis, however their role in normal and malignant hematopoiesis has not been thoroughly examined. Fibroblasts share markers with MSCs, and BM stromal cells, and lack definitive markers, thus posing a challenge in their characterization. Previously, using scRNA-sequencing, defined fibroblast clusters have been identified in normal adult BM^8^. We have similarly identified 4 fibroblast clusters in WT and preleukemic BM. We observed a drastic increase in fibroblast percentages in preleukemic BM compared to WT, as validated by imaging. Interestingly, multiple collagen genes were significantly downregulated in preleukemic BM fibroblasts compared to WT, suggesting that the BM extracellular matrix is re-modelled during the preleukemic stages.

In order to identify and further characterize the fibroblast population in preleukemic BM, we analyzed the top expressed genes and identified CD55 to be highly expressed in these cells, which was validated by fluorescence microscopy. CD55 or Decay Accelerating Factor (DAF) is a glycosylphosphatidylinositol (GPI)-anchored membrane protein that has been extensively studied as an inhibitor of complement activation^34^. CD55 has been shown to play an important role in the survival and bone-resorption activity of osteoclasts through regulation of Rac activity^35^. However, the role of CD55 and CD55+ fibroblasts in BM remains to be elucidated. This is the first study to report that CD55+ fibroblasts are increased in preleukemic BM and exhibit increased proliferation *in vitro* and *in vivo*. Gene expression analysis and microscopy reveal decreased expression of collagen genes, suggesting a remodeling of the ECM in preleukemic BM. Finally, co-culture assays suggest that CD55+ fibroblasts within preleukemic niche promote expansion of LSCs. Future studies should focus on characterizing the role of CD55 in fibroblasts and the role of CD55+ fibroblasts in leukemic progression.

## METHODS

### Single cell sequencing and data processing

Single-cell RNA sequencing (scRNAseq) was performed using 10X Genomics 3’ gene expression kit (v3.1, dual indexed). Stromal cells were sorted as previously described^8^. Briefly, WT and preleukemic mice (3 mice per group) were sacrificed, and femurs and tibias were cleaned. Bones were crushed in collagenase (Worthington Chemicals, 0.5mg/mL) and incubated at 37C for 30 mins. Cells were strained using a 70µm filter and RBCs were lysed using ACK lysis buffer on ice for 5 mins. Cells were stained with CD45, CD11b, Gr-1, B220, CD19, Ter119, CD71 and DAPI, and stromal cells were sorted by flow cytometry. Single cell suspensions of sorted stromal cells were loaded into a 10x Genomics Chromium device for microfluidic-based partitioning and 10,000 single cells were targeted for capture. The manufacturer’s instructions were followed for reverse transcription, cDNA amplification, and library preparation according to the v.3.1 3’-single-cell RNA-seq protocol. Sequencing was performed on an Illumina NovaSeq 6000 instrument to generate paired-end sequencing reads, and FASTQ files were generated using Illumina bcl2fastq software. 10x Genomics CellRanger software suite was used following the default parameters, to perform data pre-processing including alignment, filtering, barcode counting and UMI counting. Downstream analyses were performed using Seurat V3 for R^9^. For each dataset, the lowest 5% of cells based on RNA counts were discarded. Then, UMI counts were log-normalized with the NormalizeData Seurat function. For dataset integration, integration anchors were done using the FindIntegrationAnchors function. Standardized (i.e., centered and reduced) expression values were obtained with the ScaleData function of Seurat.

### Dimension reduction and clustering

Dimensionality reduction was performed by running a principal component analysis (PCA) on the scaled data using RunPCA Seurat function. The distance matrix was organized into a K-nearest neighbor graph (KNN), partitioned into clusters using Louvain algorithm using the FindClusters Seurat function, and clusters were visualized on a Uniform Manifold Approximation and Projection (UMAP) using the FindNeighbors Seurat function.

### Cluster annotation and top expressed genes

Top differentially expressed genes for each cluster were computed using the FindAllMarkers Seurat function, and cell types were annotated based on expression of known markers, as previously described^10^. Top 10 expressed genes were determined in each cluster and heatmap was generated using heatmap function.

### Differential gene expression analysis

Top differentially expressed genes in PTD; ITD stromal cells for each cluster were computed using the FindAllMarkers Seurat function. Fold changes in up- and down-regulated genes were plotted using the R package EnhancedVolcano.

### Cell cycle analysis

Cell cycle states of single cells in each cluster were identified based on expression of G2/M and S phase markers using the CellCycleScoring function in Seurat package. Percentage of cells in G1, S, and G2M cell cycle states were computed in each cluster of each group.

### Immunofluorescence

Femurs and tibias were harvested from euthanized mice and fixed in formalin for 72 hours. Bones were decalcified in EDTA solution for 72 hours. Bones were paraffin embedded and sectioned. Tumor sections were deparaffinized, sequentially hydrated and antigen retravel was performed using citrate buffer (pH 6.0). Sections were blocked and stained with primary antibodies anti-CD55 (Thermo Scientific, Cat# PA5-89530), anti-CD45 (Thermo Scientific, Cat# 14-0451-82), anti-Ter119 (Thermo Scientific, Cat# 14-5921-82), anti-Ki67 (Thermo Scientific, Cat#14-5698-82), anti-Collagen I (Thermo Scientific, Cat#MA1-26771), and anti-Collagen IV (Thermo Scientific, Cat#SAB4200500) and secondary antibodies conjugated with Alexa fluor 488, and Alexa Fluor 647. Nuclei were counterstained with DAPI. Images were obtained using the Olympus FV3000 confocal microscope and analyzed using NIH ImageJ software.

### Cell culture and MTS assay

WT and preleukemic mice (3 mice per group) were sacrificed, and femurs and tibias were cleaned. Bones were crushed in collagenase (Worthington Chemicals, 0.5mg/mL) and incubated at 37C for 30 mins. Cells were strained using a 70µm filter and RBCs were lysed using ACK lysis buffer on ice for 5 mins. Cells were stained with biotinylated lineage cocktail of CD45, CD11b, Gr-1, B220, CD19, Ter119, and CD71 for 30 minutes on ice. Cells were washed and stained with Dynabeads and incubated on shaker at 4C for 30 minutes. Cells were washed and lineage negative cells were selected using a magnetic stand. Lineage negative cells were stained with biotinylated CD55 antibody and incubated on a shaker at 4C for 40 minutes. Cells were washed and stained with Streptavidin-conjugated MACS beads at 4C for 15 minutes followed by positive selection through MACS LS column. CD55+ cells were cultured in RPMI supplemented with 20% FBS, and antibiotics. Media was changed every 3-4 days and cells were passaged every 7-10 days. P2 CD55+ cells were plated in a 96 well plate and MTS assay was performed according to manufacturer’s instructions at 24, 48, and 72 hours post plating. For co-culture assay, cKit+ cells were isolated from WT and pre-leukemic PTD;ITD mice BM using MACS cKit isolation kit according to manufacturer’s protocol, and co-cultured with CD55+ cells from WT or pre-leukemic PTD;ITD mice for 48 hours, followed by colony formation assay as previously described^11^. Colonies were counted 14 days later.

