## Supplemental Data for "Cellular taxonomy of the preleukemic bone marrow niche of acute myeloid leukemia"

#### SUPPLEMENTAL FIGURES

#### Supplemental Figure S1. Flow cytometry gating strategy for sorting

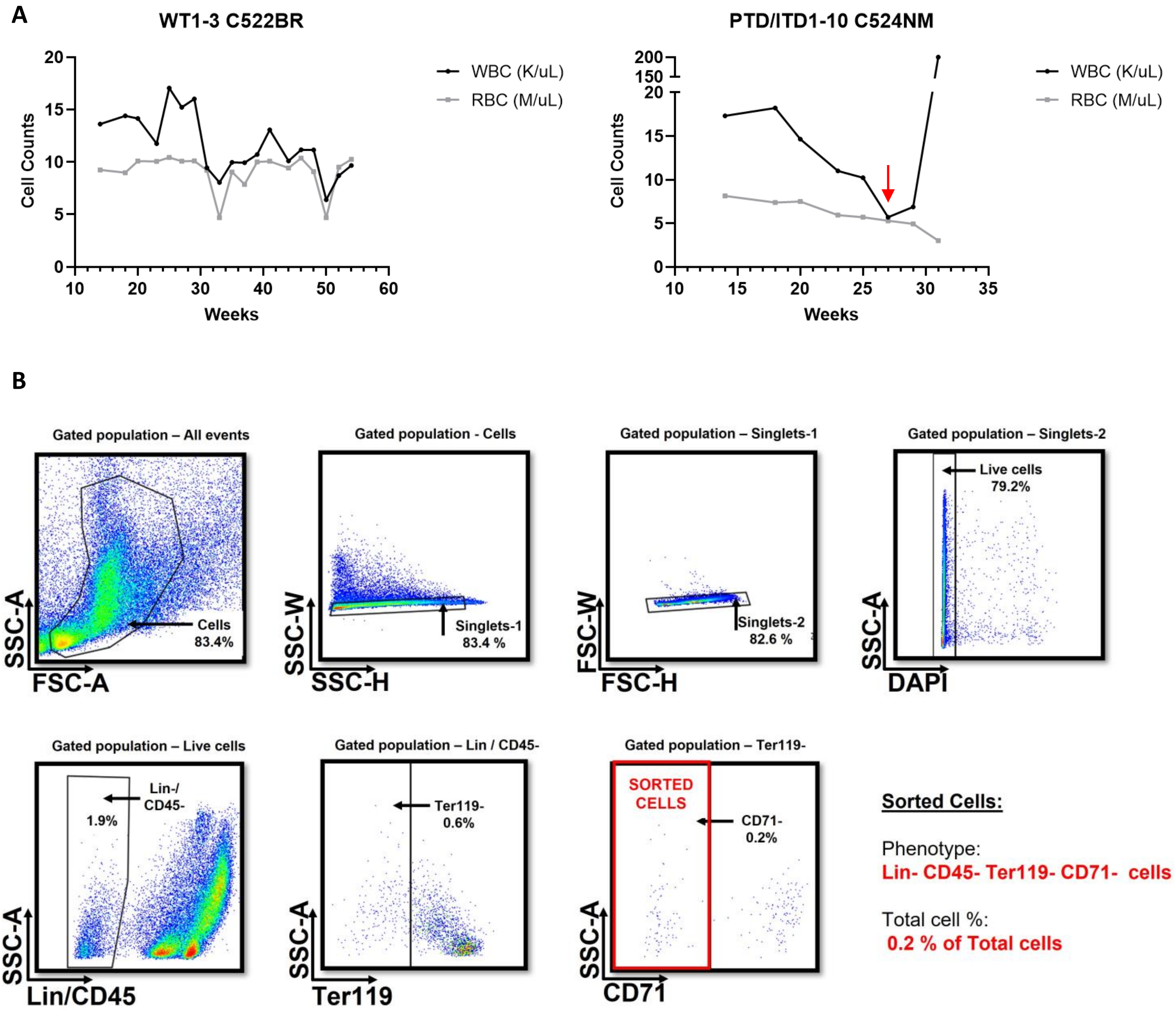


**Supplemental Figure S1.** Isolation of stromal cells from preleukemic PTD; ITD mice. (A) Peripheral blood WBC, and RBC counts from WT and PTD; ITD mice over time (weeks). Mice were bled every 2-3 weeks and WBC, and RBC counts were taken to determine 'preleukemic' stage. (B) Representative gating strategy to sort CD45- Lin- (CD3- B220- CD11b- Gr1- CD19-) Ter119- CD71- stromal cells.

**Supplemental Figure S2. Proliferation profiles of BM stromal cells**

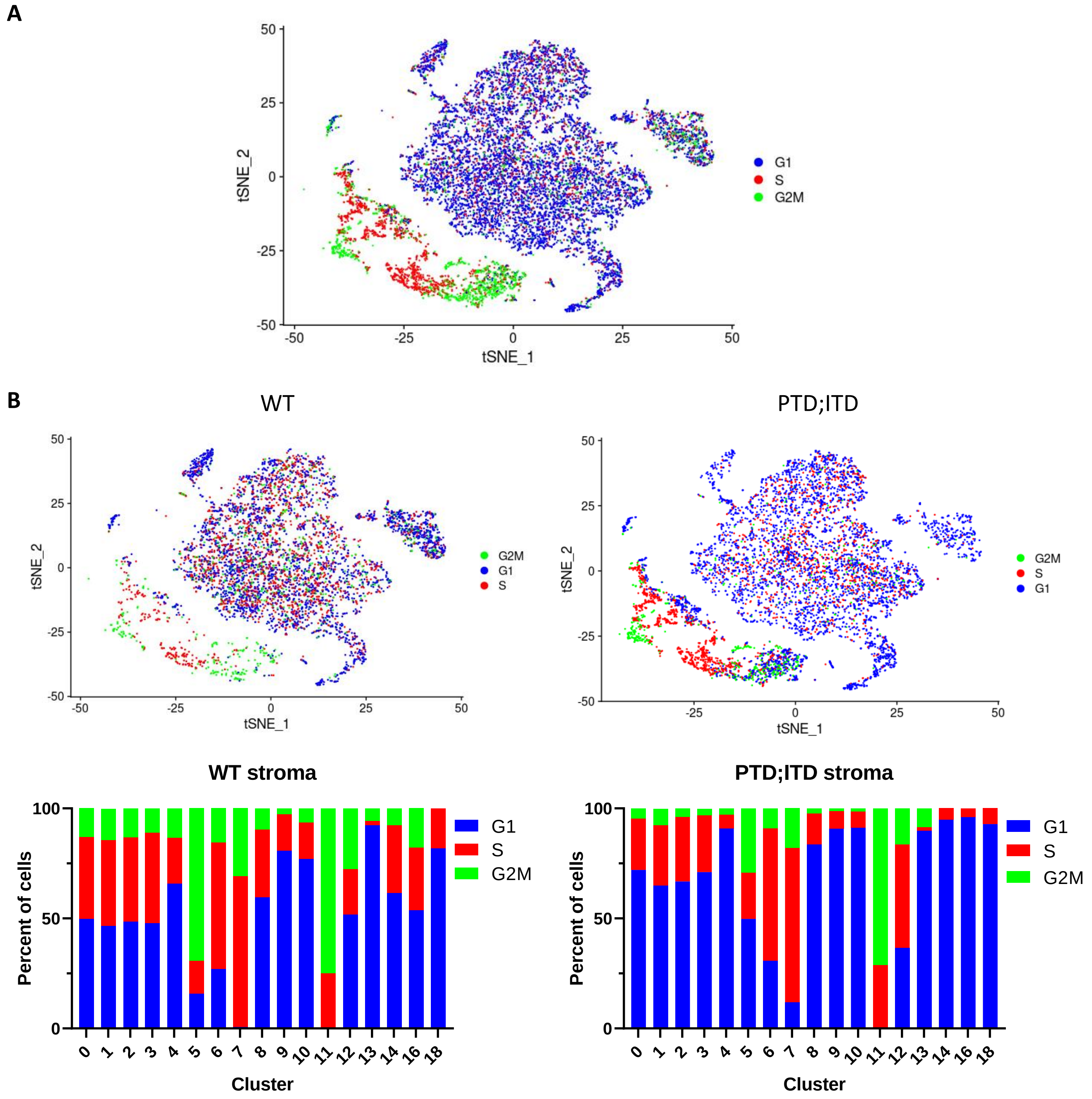


**Supplemental Figure S2.** Proliferation profiles of BM stromal cells. (A) t-SNE of stromal cells by cell cycle phases G1, S, and G2M. (B) (top) t-SNE of stromal cells from WT and preleukemic PTD; ITD mice by cell cycle phases G1, S, and G2M. (bottom) Percentage of stromal cells in G1, S, and G2M phases in each cluster of WT and PTD; ITD stromal cells.

### Supplemental Figure S3. Genes expressed in LepR MSC clusters

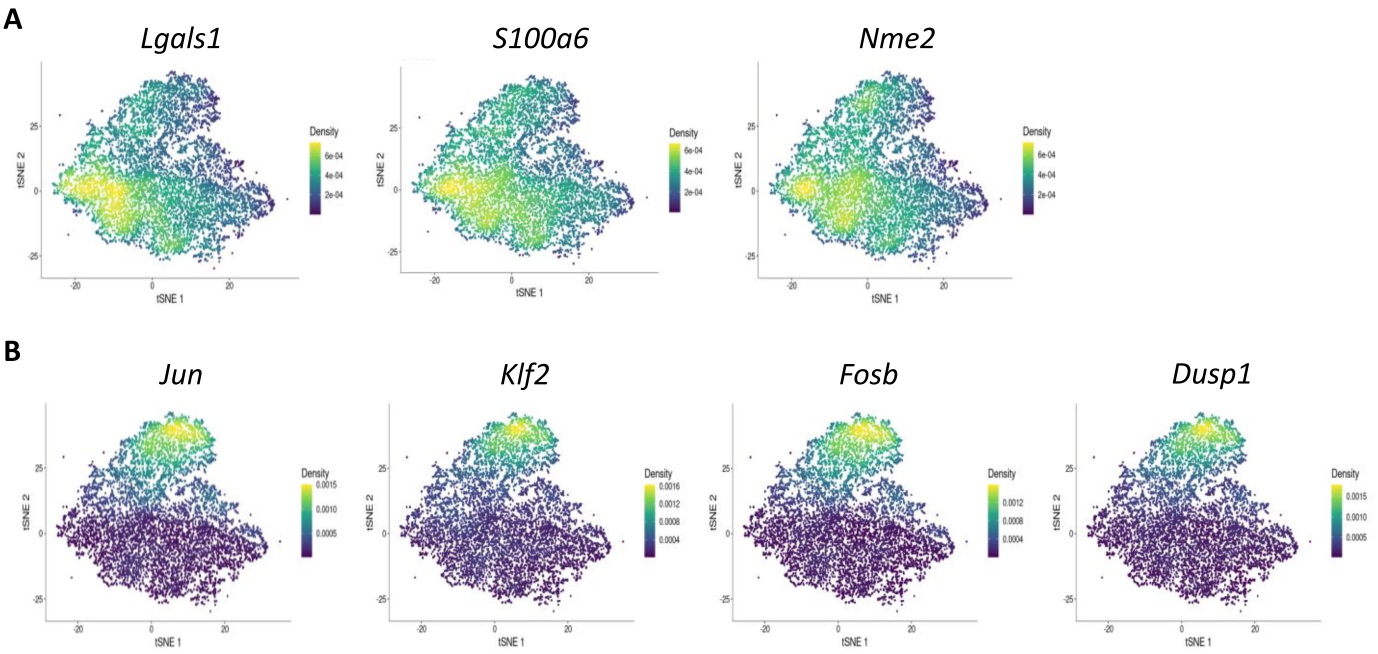


**Supplemental Figure S3.** Genes expressed in LepR MSC clusters. t-SNE of LepR MSCs by expression of (A) genes differentially expressed in cluster 0, and (B) genes expressed in stem-like MSCs.

#### Supplemental Figure S4. Osteo-lineage cells

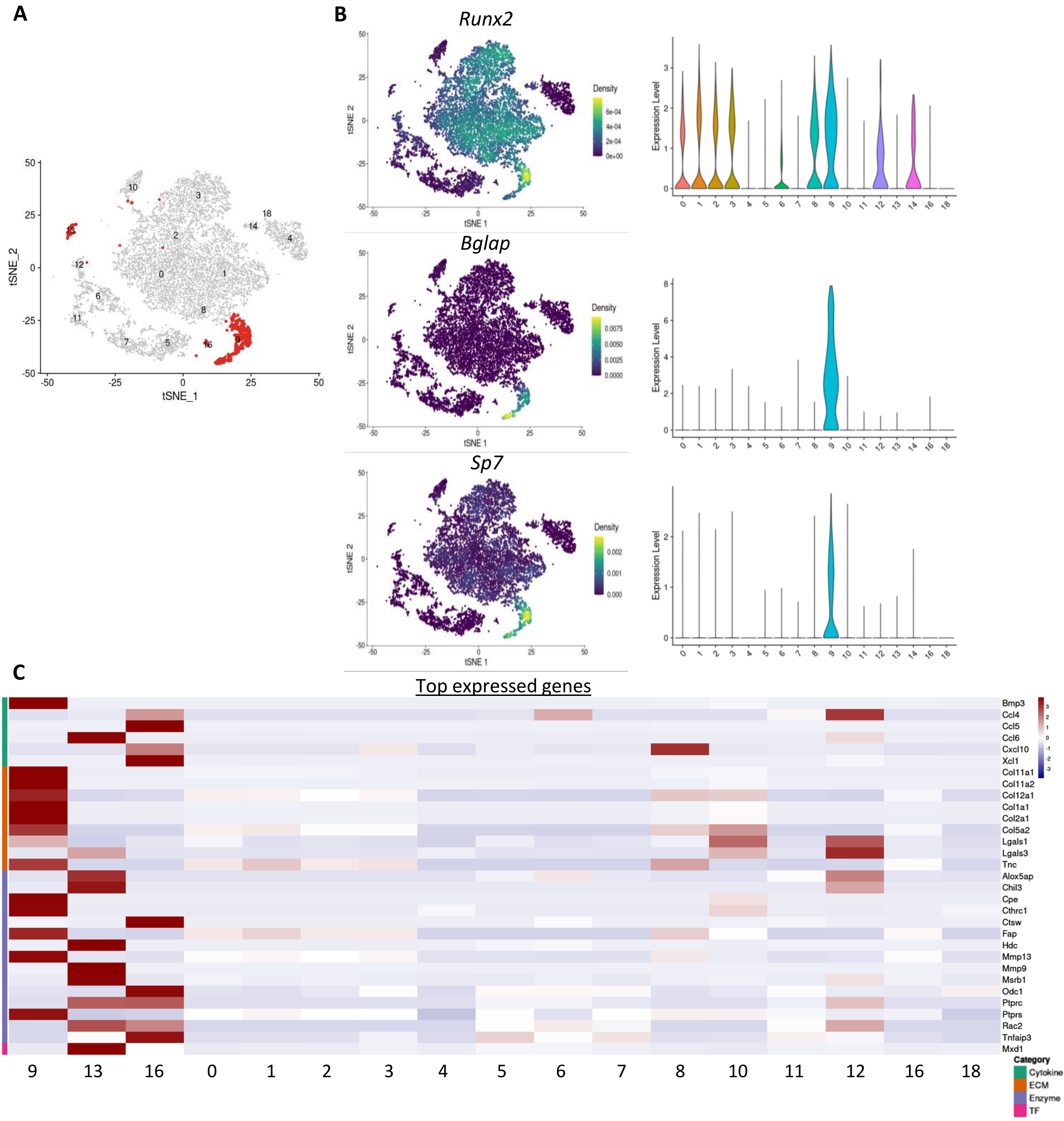


**Supplemental Figure S4.** Osteo-lineage cells. (A) t-SNE of stromal cells, highlighting osteo-lineage cell clusters. (B) t-SNE of stromal cells by expression of key osteo-lineage marker genes, and corresponding violin plot distributions of expression levels across clusters. (C) Expression of top expressed genes (rows) of osteo-lineage cells in the cells of each cluster (columns) ordered by gene categories.

#### Supplemental Figure S5. Endothelial cell clusters

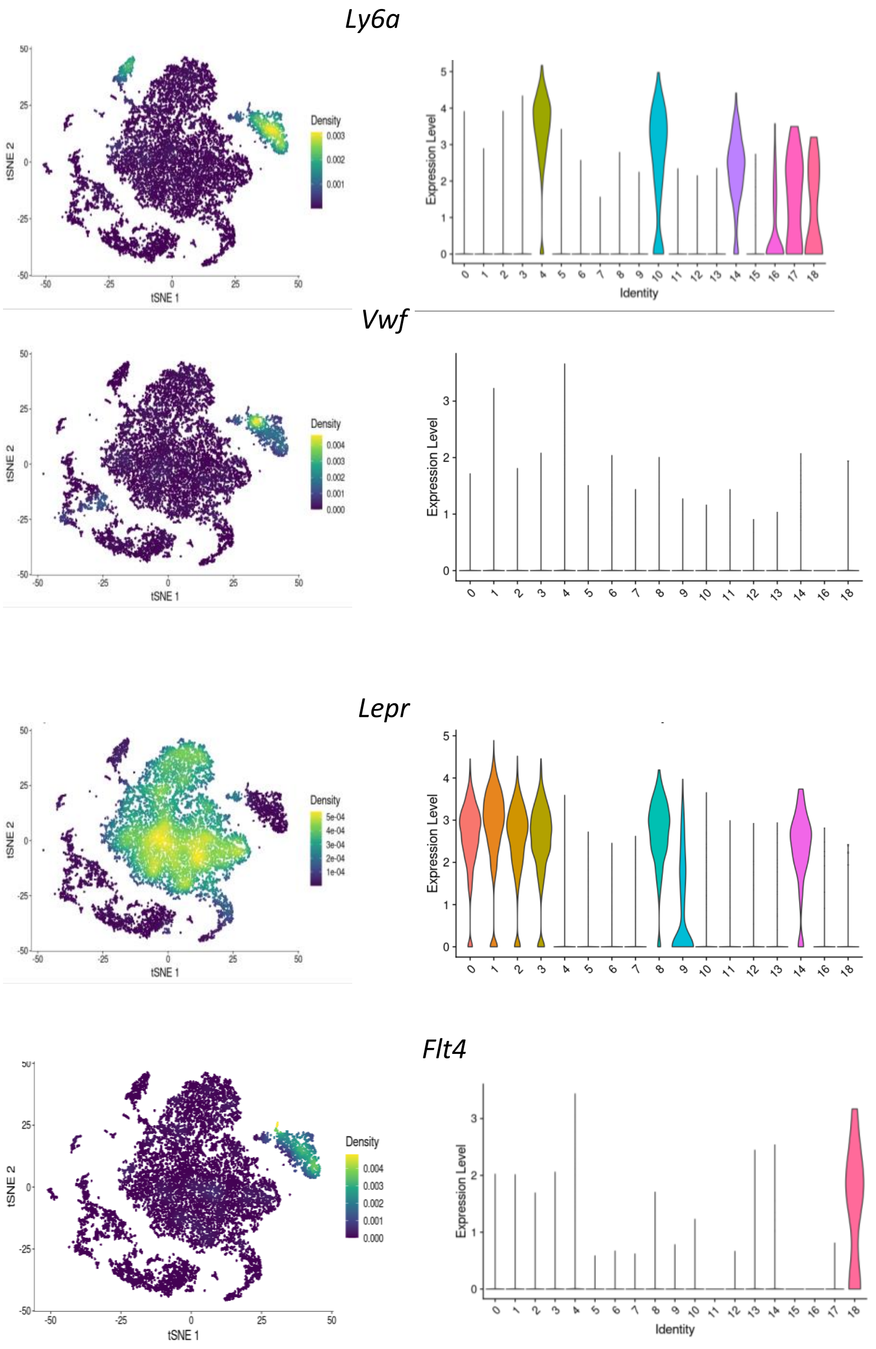


**Supplemental Figure S5.** Endothelial cell clusters. t-SNE of stromal cells by expression of endothelial cell-specific markers, and corresponding violin plot distributions of expression levels across clusters.

### Supplemental Figure S6. Genes expressed in fibroblast clusters

**A** Transcription factors

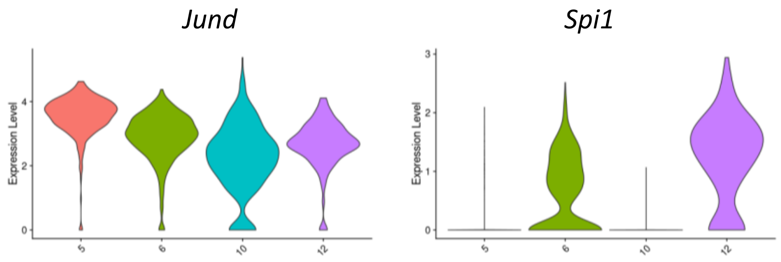


**B** Cell surface markers

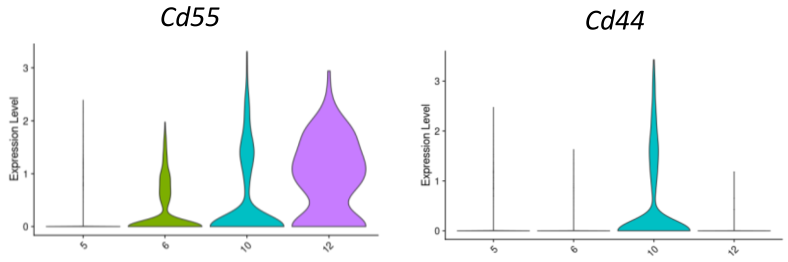


**Supplemental Figure S6.** Genes expressed in fibroblast clusters. (A) Violin plots of expression of transcription factor genes across fibroblast clusters. (B) Violin plots of expression of cell surface marker genes across fibroblast clusters.

#### Supplemental Figure S7. scRNA-seq analysis of sorted CD55+ fibroblasts

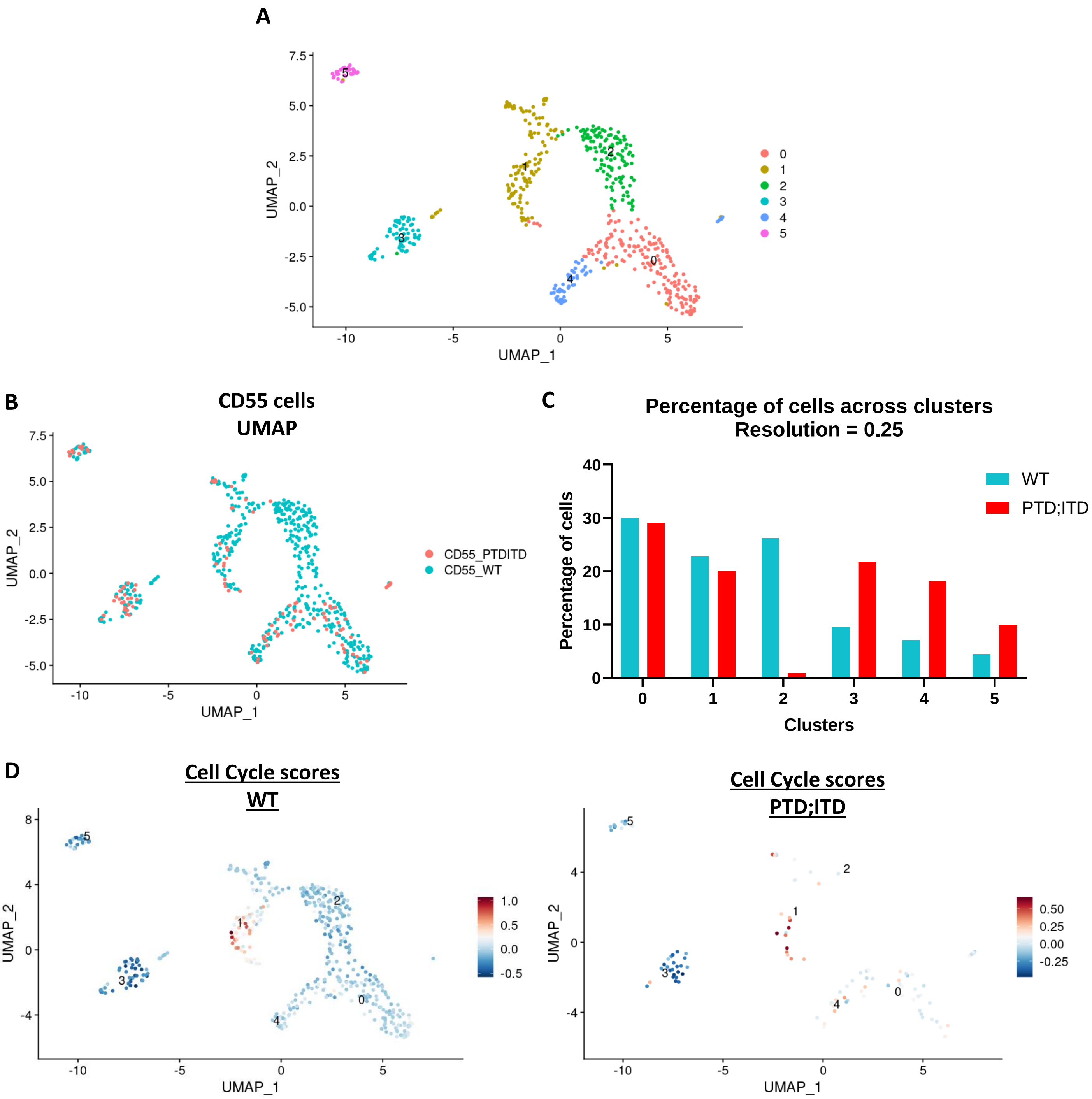


**Supplemental Figure S7.** scRNA-seq analysis of sorted CD55+ fibroblasts. (A) Uniform manifold approximation and projection for dimension reduction (UMAP) of sorted CD55+ stromal cells, colored by clustering. (B) UMAP of CD55+ stromal cells colored by condition (WT, blue; preleukemic PTD;ITD, red). (C) Percentage of CD55+ stromal cells across each cluster in WT and preleukemic PTD; ITD cells. (D) UMAP of cell cycle scores in WT (left) and PTD;ITD (right) CD55+ stromal cells.
